# Male youth BMI is associated with epigenetic ageing of female descendants but not blood or sperm DNA methylation

**DOI:** 10.64898/2026.10.26.761759

**Authors:** Jo Ciantar, Saara Marttila, Ilari Taskinen, Virva Liski, Noora Kartiosuo, Damien Kaukonen, Pashupati P. Mishra, Leo-Pekka Lyytikäinen, Suvi Rovio, Juha Mykkänen, Katja Pahkala, Mika Kähönen, Noora Kotaja, Terho Lehtimäki, Olli Raitakari, Ville Kivimäki, Emma Raitoharju

## Abstract

Studies based on harvest records indicate that grandparental nutrition in youth is associated with the health traits of grandchildren in a sex-specific lineage. We investigated whether male early adulthood anthropometrics associate with epigenetic age, a measure of general well-being, and DNA methylation, a potential molecular inheritance mechanism, in three generations of descendants.

Data on weight, height, BMI and chest circumference measured at military draft examinations (at age 18–24 yrs.) were retrieved from national archives for 1,013 men, herein referred to as Generation Archives (G_A_). The genome-wide DNA methylation of their descendants participating in the Young Finns Study (YFS) was measured from the blood and sperm from the children (59–80 yrs., n_blood_ = 362, n_sperm_ = 51), grandchildren (34–49 yrs., n_blood_ = 421, n_sperm_ = 98) and great-grandchildren (7–30 yrs., n_blood_ = 200).

The men’s (G_A_) BMI had an inverse association with the epigenetic pace of ageing (DunedinPACE) of their daughters (β = −0.01, p = 0.002, n = 169) and the daughters’ daughters (β = −0.01, p = 0.049, n = 126) but not with that of their male descendants. From within the YFS cohort, maternal, but not paternal, BMI at ages 13–34 also had an inverse association with the DunedinPACE of the immediate offspring. However, robust evidence for the inheritance mechanism was not found in the analyses between the anthropometric measurements of the G_A_ men and the DNA methylation profiles in the blood and sperm of their descendants.

## Introduction

Experiments on rodents have shown transgenerational inheritance of acquired traits in mammals. In particular, paternal dietary changes have been shown to affect the metabolic traits of the following generations [1]. Some of the effects have been shown to last for up to four generations [2,3]; some dilute over time, some are specific to maternal or paternal lineage, while others present solely or predominantly in male or female offspring. In mice, spermatozoa RNAs have been shown to mediate the epigenetic inheritance of acquired traits [4–7] and influence the early embryonic development, including the establishment of the epigenomic profile of future offspring [8]. Additionally, studies have found that the methylation state of some regions that are inherited by offspring are also preserved in mature sperm [9]. Man-made modifications of DNA methylation have also been shown to be passed on to offspring in mice [10].

DNA methylation profiles are mostly erased during the development of primordial germ cells and again in the early embryo [11]; therefore, they have often not been considered to be mediators of epigenetic inheritance. However, up to 5,000 loci in the human genome have been shown to escape the first [12] and approximately 100 genes the second, demethylation wave [13]. Studies on epigenetic inheritance have primarily searched for replicative transmission of epigenetic marks and dismissed the possibility of reconstructive transmission, in which the primary epigenetic marks, i.e. DNA methylation, are erased in gametogenesis and then faithfully reconstructed after fertilization via the inheritance of a secondary carrier, such as ncRNAs [14].

Human studies on intergenerational inheritance of acquired traits where the parental exposure has been inflicted before the pre- or periconceptional period remain sparse. Parental weight status, psychological stress and exposure to toxins and tobacco smoke prior to the pre-/periconceptional period have been associated with the birthweight, body weight, psychosocial traits and respiratory health of offspring [15]. In addition, the parental age at the birth of the offspring has been linked to the risk of autism [16,17] and to changes in epigenetic profiles [18,19].

For an effect to be truly considered transgenerational, timing of the exposure would need to take place before development of gametes of the future offspring [1,20]. Therefore, all grandpaternal exposures as well as exposures happening in a grandmother who is not pregnant are considered transgenerational. Human studies show that the paternal grandfather’s access to food in pre-puberty predicts the all-cause-mortality of his male, but not female, grandchildren [21,22] and food availability during the youth of the maternal grandmother has been associated with the mental health of her daughters’ daughters [23]. In addition to lineage specificity, a study by Sari et al. indicates that *in utero* exposure may have contradictory effects depending on the socioeconomic status. The same negative exposure in mothers with economic well-being may leads to “scarring” of their descendant’s health, while in mothers experiencing economic hardship, it may lead to culling, and surviving descendants may thus show improved health [24]. These studies suggest that transgenerational inheritance or repeated intergenerational inheritance of acquired traits also exists in humans. Very little is known about the mechanisms conveying it, with few studies indicating changes in blood DNA methylation patterns in descendants of those exposed to war-related violence [25] or the Shoah (Holocaust) [26]. There are currently no studies investigating potential transgenerational effects on the sperm DNA methylation of descendants in humans.

Inspired by the studies based on historical harvest records, we set out to examine whether the anthropometric measures (height, weight, BMI and chest circumference) of Finnish men, as measured at military draft examinations (in 1927–1957, at ages 18–24), were associated with the blood and sperm DNA methylation patterns of their children, grandchildren, or great-grandchildren [27]. Furthermore, we aimed to investigate whether the anthropometric measurements associated with the overall health status of descendants, as measured with epigenetic clocks computed from blood DNA methylation data. The included epigenetic clocks represent lifespan and health span (PhenoAge [28]), the risk of morbidity and mortality (GrimAge [29]) and the pace of biological ageing (DunedinPACE [30]). In addition, we utilized the three-generational Young Finns Study (YFS) cohort [27] to study whether the findings can be replicated between parental anthropometrics and the children’s epigenetic pace of ageing.

## Results

### Study Population

Anthropometric data of 994–1,013 men, recorded in military draft records when they were under the age of 25, were obtained from the National Archives of Finland. These men, referred to as the Generation Archives (G_A_), were linked to three generations of their descendants in the YFS cohort [27]. The children, grandchildren and great-grandchildren of the G_A_ men are referred to as G0, G1 and G2, respectively. Longitudinal data from 1980–2020 are available for the original YFS participants (G1 – the grandchildren of the G_A_ men), while data on the other two generations were only collected during the 2018–2020 follow-up [27] (Figure 1).

**Figure 1:**
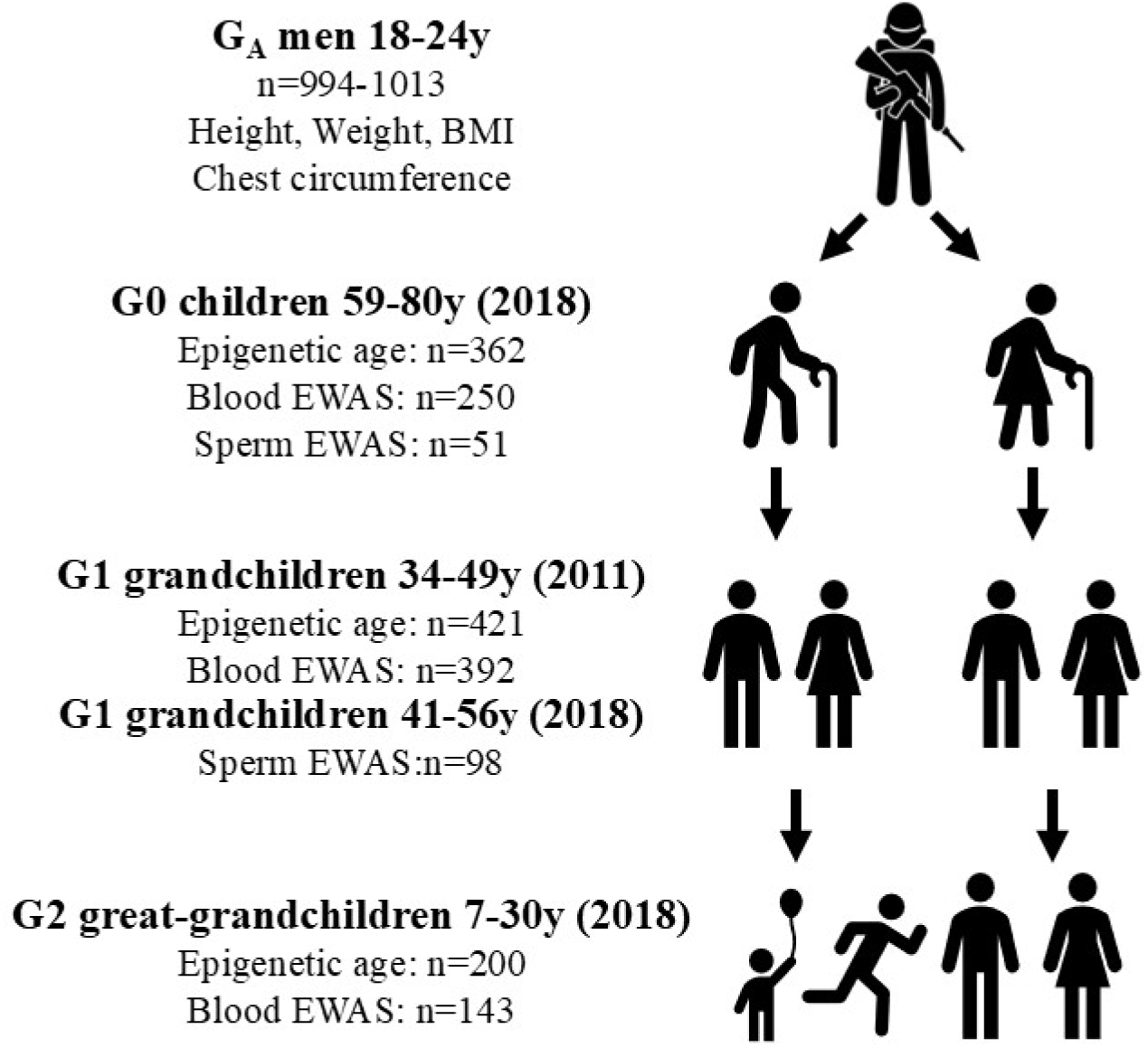
Study population Our study cohort consists of four generations: the G_A_ men for whom anthropometric data are available, as well as their children (G0), grandchildren (G1) and great-grandchildren (G2), for whom DNA methylation data are available. For G1, blood methylation data that were collected in the 2011 follow-up were used, as this follow-up contained the largest number of individuals. Sperm methylation data were only collected in the 2018–2020 follow-up. Data for G0 and G2 were only collected in the 2018–2020 follow-up. For some of the G1 participants, data are available for both maternal and paternal G_A_ grandfathers (i.e. their mother’s father and their father’s father), so the sum of individuals included in the lineage-specific analysis is more than the number of participants shown in this figure. When data from multiple siblings from G2 were available, only one was retained in the dataset by random selection to maintain variable independence.

The height, weight, chest circumference and BMI of the G_A_ men were all normally distributed (Supplementary Figure 1). A BMI of under 18.5 was observed in 2.2% of the men, and 3.1% had a BMI of over 25. Two individuals had a BMI higher than 30 and were thus considered to be outliers and removed from downstream analyses. The height of 6.1% of the G_A_ men was under 160 cm, and 11% weighed less than 55 kg (Table 1). All anthropometric measures except BMI correlated with the year of birth, but none correlated with age at data measurement. Only height and weight were correlated with social class, with members of the upper class being both taller and heavier than those of the lower classes (Supplementary Table 1).

**Table 1:**
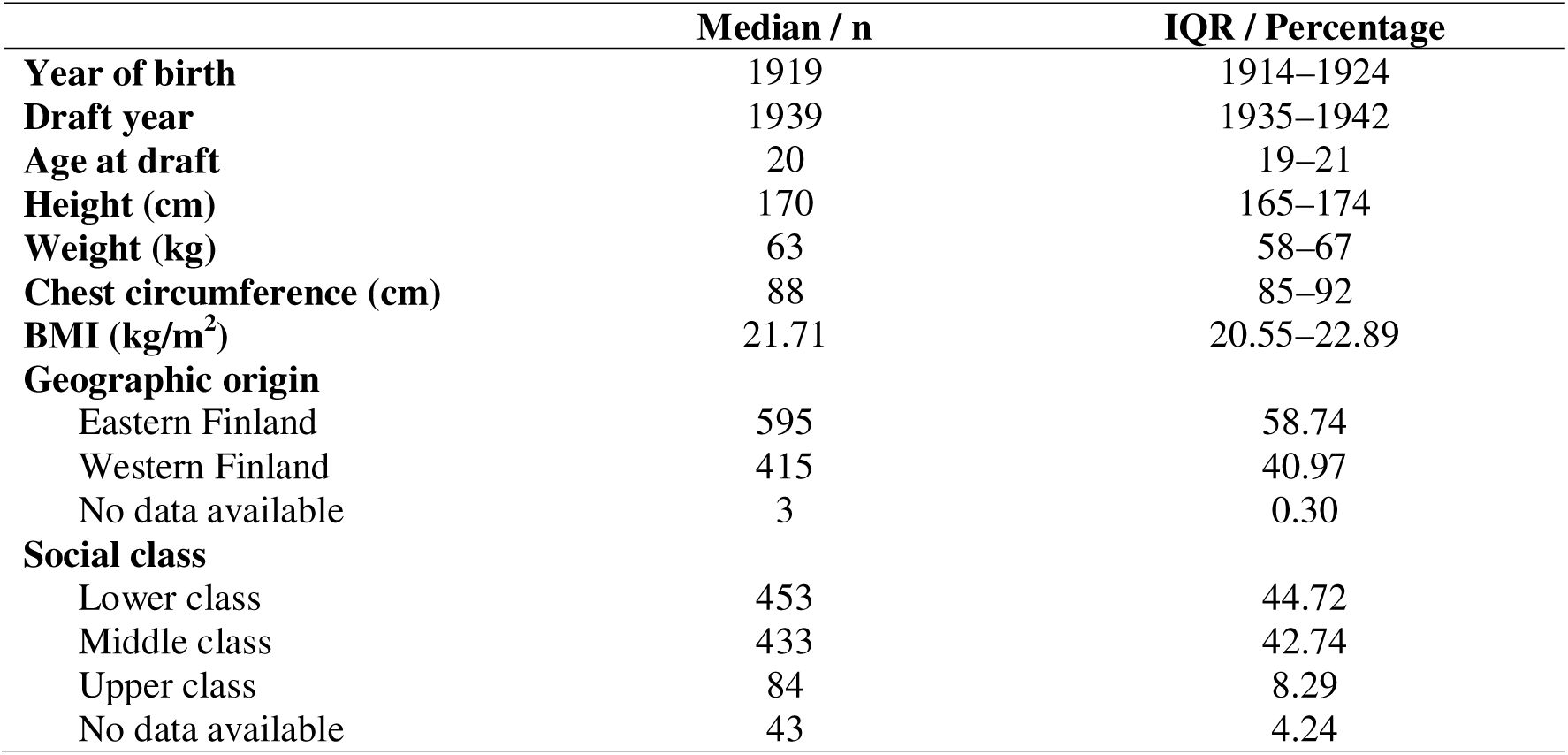
Demographics of the G_A_ men The year of birth, height, weight, chest circumference and age of the G_A_ men were obtained from military draft records, and the BMI was calculated. Geographic origin was categorized from birth municipality data, and social class categories are based on the recorded occupation. Medians and interquartile ranges (IQR) are given for continuous variables, while the number of individuals and percentages are given for categorical variables. Demographics of the descendants can be found in Supplementary Table 2.

### Association of the G_A_ men’s anthropometric measurements with epigenetic ageing in three subsequent generations (G0, G1 and G2)

The association between each anthropometric trait of the G_A_ men and each epigenetic age measure in three generations of their descendants was investigated using three regression models (Figure 2). The basic model was only adjusted for the descendants’ (G0/G1/G2) features, while the adjusted model was additionally adjusted for the G_A_ men’s features, and the fully adjusted model also included the years of schooling of the descendants. Stratification by sex and/or lineage was also performed, when appropriate. The investigated epigenetic age measures were the age deviation (AgeDev) values of the principal-component versions of PhenoAge and GrimAge, as well as DunedinPACE.

**Figure 2:**
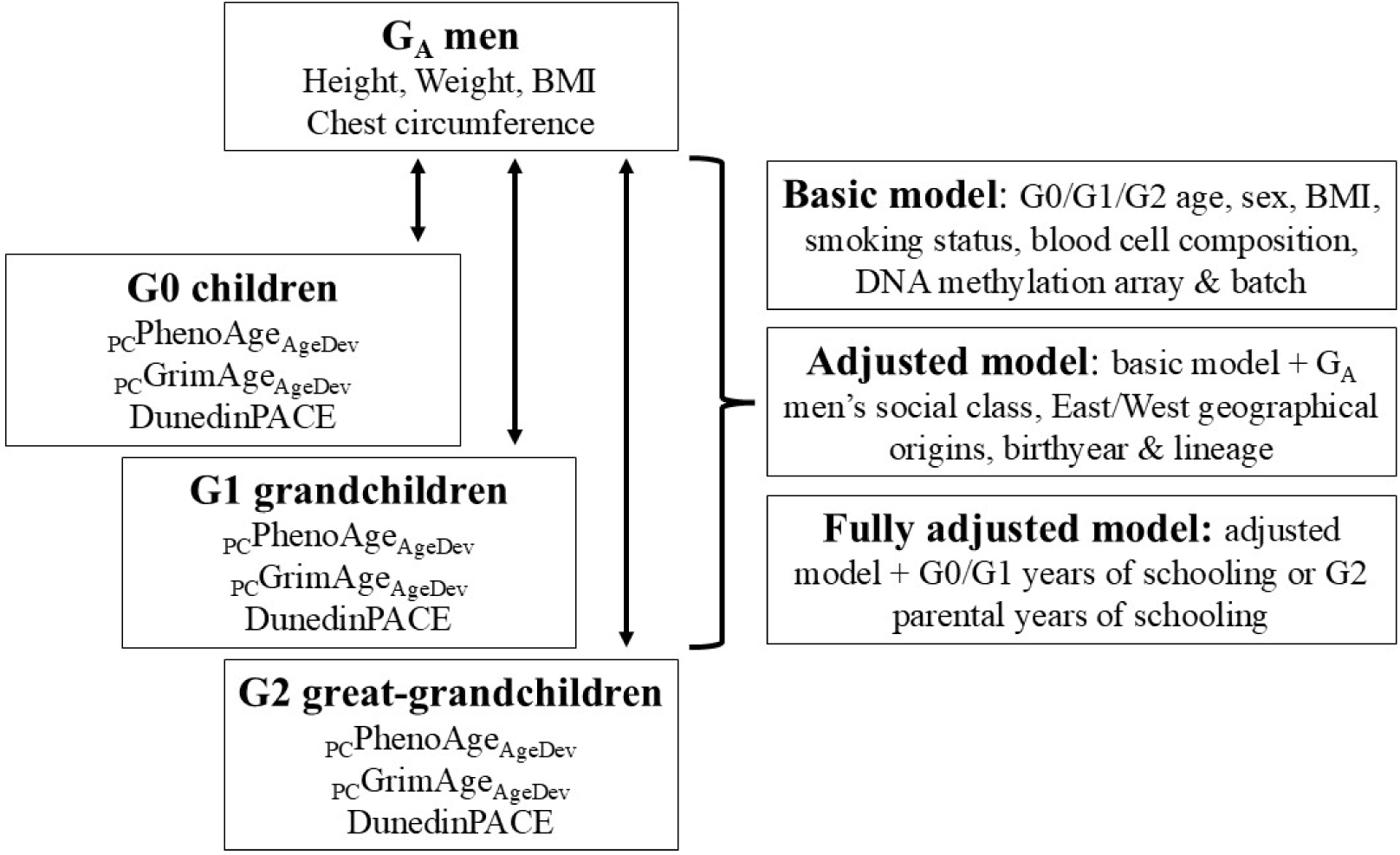
Analyses schematic Three models were used to investigate the association between each anthropometric trait of the G_A_ men and each epigenetic age measure of three generations of their descendants, separately. The basic model was adjusted for features of the descendant that are known to associate with epigenetic age, the adjusted model was additionally adjusted for features of the G_A_ men, and the fully adjusted model also included the years of schooling for G0 and G1. Stratification by sex and/or lineage, where appropriate, was also performed.

A systematic association was seen between the G_A_ men’s BMI at drafting and their descendants’ DunedinPACE, with a higher BMI being associated with a slower pace of epigenetic ageing (Figure 3 and Supplementary Tables 3–5). Inverse associations were observed in all three models with all children (β = −0.008, p = 0.010–0.017), daughters (β = −0.01, p = 0.002–0.006) and maternal grandchildren, i.e. the daughters’ children (G1; β= −0.007–-0.008, p = 0.010–0.023). Additional inverse associations were also observed, albeit only in the basic and adjusted models, in maternal granddaughters, i.e. daughter’s daughters (G1; β = −0.01, p = 0.015–0.049). While no significant associations were observed between G_A_ BMI and descendant _PC_PhenoAge_AgeDev_, we also found inverse associations between the G_A_ men’s early adulthood BMI and _PC_GrimAge_AgeDev_ in the basic models with maternal grandchildren and maternal granddaughters (β = −0.17–-0.23, p = 0.030–0.048).

**Figure 3:**
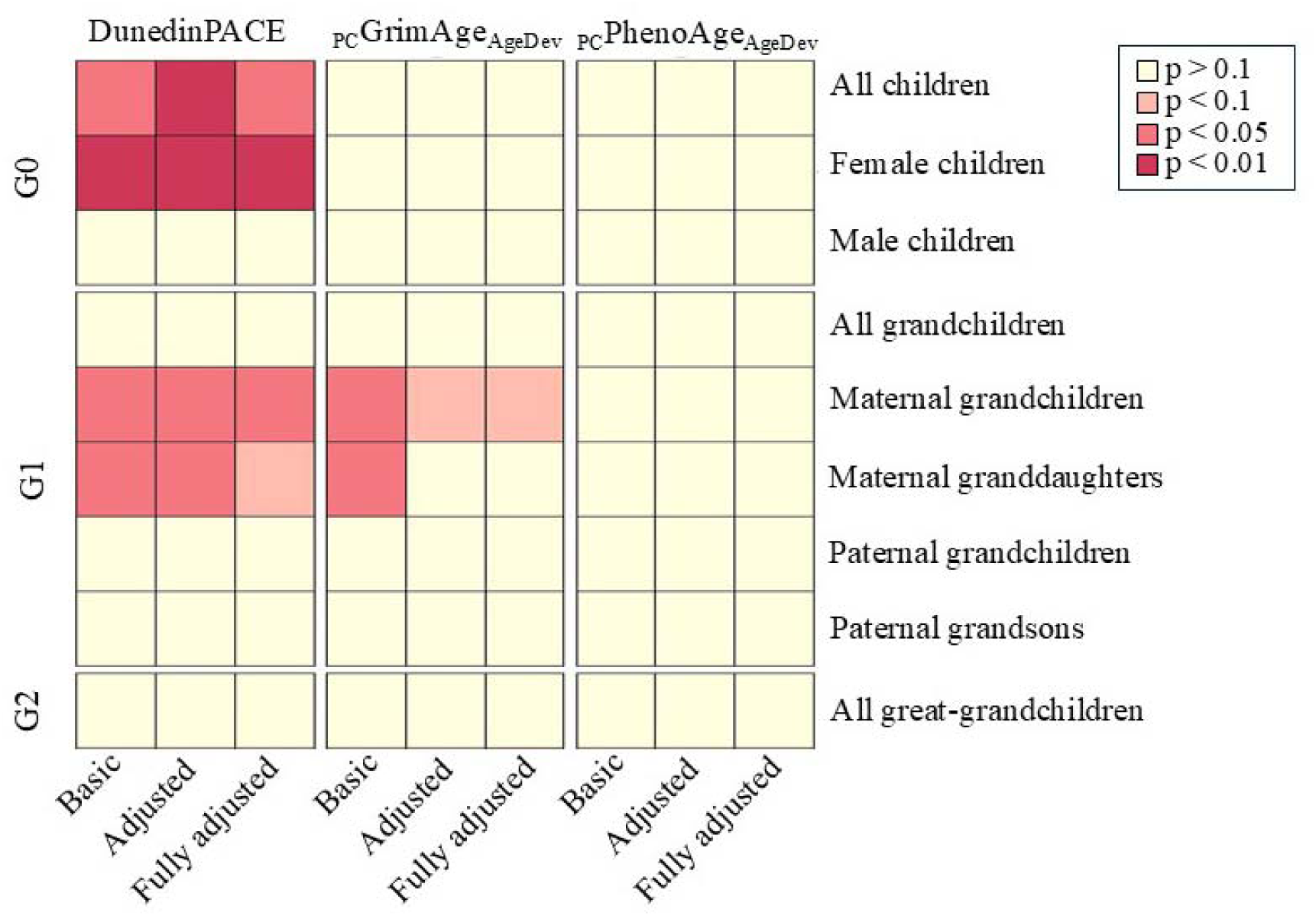
The association between G_A_ men’s BMI and their descendants’ epigenetic age The BMI of G_A_ men shows a consistent inverse association with their descendants’ pace of epigenetic ageing, especially in the maternal line. For each regression model, the early adulthood BMI value of the G_A_ man was treated as the independent variable, while the epigenetic age measure of the descendant was treated as the dependent variable. The basic model was adjusted for the descendant’s age, sex, BMI, smoking status, methylation-based estimates of cell proportions, as well as the DNA methylation array and batch, when applicable. The adjusted model was further adjusted for the G_A_ men’s social class, geographic origin, year of birth and lineage, when applicable. The fully adjusted model was additionally adjusted for the years of schooling that the descendant or, in the case of G2, their parent had completed. All associations are inverse and the colour intensity corresponds to the p value.

As a sensitivity analysis, we repeated the analysis for female children, maternal grandchildren and maternal granddaughters and included only participants who did not smoke. The inverse associations replicated in the non-smoker subset and the fully adjusted model for maternal granddaughters also reached the threshold of statistical significance (β = −0.013, p = 0.02; Supplementary Table 6).

We then categorized the BMI values into two groups to compare the individuals in the lowest or highest quartile to the rest. Significant associations were only seen in the comparison of the highest BMI quartile (BMI > 23), suggesting that the association is primarily driven by the individuals in the upper quartile. Daughters of the G_A_ men who were in the highest BMI quartile aged a little over two weeks less per year, as measured by DunedinPACE, than the daughters of the men (G_A_) in the lower BMI group (basic model β = −0.05, p = 0.002, n = 178). The magnitude was similar for the maternal granddaughters (basic model β = −0.041, p = 0.033, n = 132; Supplementary Table 7).

Some spurious associations were also found for the other anthropometric measurements, but not as systematically as in the BMI analysis. The early adulthood height of the G_A_ men had a direct association with their grandchildren’s _PC_PhenoAge_AgeDev_ and DunedinPACE in the adjusted and fully adjusted models (β = 0.001–0.039, p = 0.031–0.049) but not in the sex-stratified analyses (Supplementary Figure 2 and Supplementary Tables 8–10). The weight of the G_A_ men had an inverse association with the maternal grandchildren’s and maternal granddaughters’ DunedinPACE, and with the maternal granddaughters’ _PC_GrimAge_AgeDev_ in the basic model (β = −0.003–-0.058, p = 0.033–0.048; Supplementary Figure 3, Supplementary Tables 11–13). Chest circumference did not associate with any of the descendants’ epigenetic age measures at p < 0.05 (Supplementary Figure 4 and Supplementary Tables 14–16).

### Internal replication in association between parental early-life BMI and the offspring’s DunedinPACE

We next sought to replicate some of our findings by investigating whether the early-life BMI of the original YFS participants (G1), for whom longitudinal data are available, was associated with the DunedinPACE of their offspring (G2). BMI measurements from 1980 to 2011 were used to calculate the area under the curve (AUC) and to extrapolate estimate values for each year between the ages of 3 and 34, as described previously [31]. Regression models were applied adjusting with G2 age, smoking status and BMI; G1 age, geographic origin and years of schooling; as well as DNA methylation batch, array version and cell proportions. Analyses were performed separately for the mothers and fathers of all children and then stratified by the sex of the offspring. When data of more than one G2 offspring from the same G1 parent were available, a sibling was selected at random to be included in the analysis to maintain variable independence.

The mothers’ estimated BMI from age 13 upwards was inversely associated with the DunedinPACE of their children (Figure 4A; Supplementary Table 17). In the sex-stratified analyses, a similar statistically significant inverse association was observed for sons with estimates of maternal BMI at 21 years and older. In daughters, the association was not statistically significant at any maternal age. No associations were identified between the fathers’ BMI and the DunedinPACE of their offspring (Figure 4B; Supplementary Table 18).

**Figure 4:**
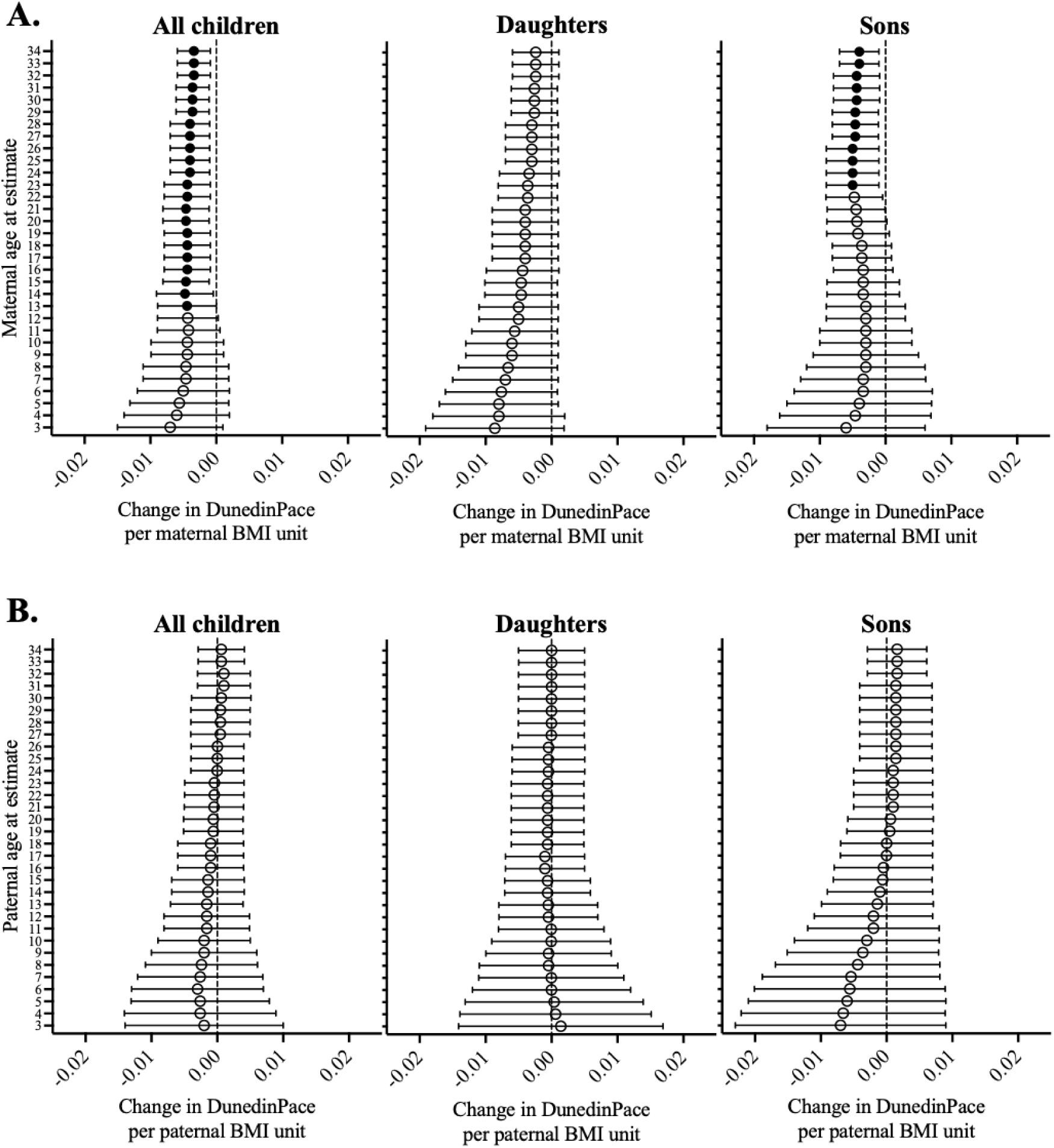
The estimated **A.**) maternal (n = 255) and **B.**) paternal (n = 191) BMI between the ages of 3 and 34 in association with the DunedinPACE of the offspring. In regression analyses, paternal/maternal (G1) BMI was treated as the independent variable and the DunedinPACE of the offspring (G2) as the dependent variable. Models were adjusted for offspring (G2) age, sex, BMI and smoking status; parental (G1) age, geographic origin and years of schooling; as well as methylation batch, array version and estimated cell proportions. When data from multiple offspring for the same parent were available, one *sibling was selected at random to be included in the analysis. Associations with a nominal p-value of < 0.05 are indicated by a full circle*.

### G_A_ men’s anthropometric measurements and the genome-wide blood DNA methylation of their descendants

Epigenome-wide association studies (EWASs) were performed to determine whether the G_A_ men’s anthropometrics associate with their descendants’ DNA methylation at each of the loci measured by the Illumina arrays. Analyses of the children (G0) were performed all together (n = 250) and separately for sons (n = 133) and daughters (n = 117), while the grandchildren (G1) were analysed all together (n = 392) as well as separately for paternal grandchildren (n = 242), maternal grandchildren (n = 253), paternal grandsons (n = 100) and maternal granddaughters (n = 121). Great-grandchildren (G2) were analysed all together (n = 143), as stratification would have resulted in too small sample sizes. All analyses were adjusted for the G_A_ man’s social class, geographic origin and year of birth, as well as the descendant’s age, BMI and smoking status, in addition to the blood cell composition and the first 10 principal components of the methylation control probes. Where appropriate, the models were also adjusted for lineage and the descendant’s sex.

No statistically significant (FDR < 0.05) associations were found between the G_A_ men’s anthropometrics and their children’s (G0) blood DNA methylation pattern when analysed all together or when stratified by sex (associations with nominal p-value of < 0.0001 can be found in Supplementary Tables 19–30). Similarly, almost no statistically significant associations were found between the G_A_ men’s anthropometrics and their grandchildren’s (G1) DNA methylation patterns when analysed either together, stratified by linage or stratified by lineage and sex (nominal p < 0.0001 associations in Supplementary Tables 31–50). The only exception was between the G_A_ men’s BMI and maternal grandchildren’s DNA methylation at cg10866062 in the transcription start site of *BTBD17* (FDR = 0.03, β = 0.008). In great-grandchildren (G2), no statistically significant associations were found (FDR > 0.05; associations with a nominal p-value of < 0.0001 can be found in Supplementary Tables 51–54).

### G_A_ men’s anthropometric measurements and the genome-wide sperm DNA methylation of their sons (G0) and grandsons (G1)

DNA methylation from purified sperm of sons (G0) and grandsons (G1) was measured with reduced representation bisulphite sequencing (RRBS) for a subset of the study cohort. The analyses for sons (G0, n = 51) and grandsons (G1, n = 98) were performed separately, and the models were adjusted similarly to the blood analyses. The analysis for great-grandsons (G2) could not be performed, as data were available for too few individuals (n = 24).

In both sons (G0) and grandsons (G1), one statistically significant differentially methylated locus was identified (FDR < 0.05). In sons and grandsons, increased methylation in chr2:231703705 and chr17:22506442, respectively, was associated with an increasing early-adulthood BMI of the G_A_ men. (Supplementary Tables 55 and 56).

## Discussion

When investigating the association of the anthropometrics of men (G_A_) in adolescence and young adulthood with measurements of epigenetic ageing in their descendants, we found a systematic inverse association between the men’s BMI and the DunedinPACE of their daughters and the daughters’ children. A similar trend was observed between the body weight of these men and the DunedinPACE of their daughters’ daughters. While the effect of BMI on the descendants’ pace of ageing was driven by the highest BMI quartile (BMI > 23), the majority of the cohort had a normal BMI (95% had a BMI between 18.5 and 25), indicating that it is the higher end of the normal range of BMI that was beneficial to the epigenetic ageing of the descendants in our cohort, not overweight or obesity.

Previous literature has linked grandparental nutrition availability with the health of the descendants, and famine, in particular, has been associated with the health of the grandchildren [21–23]. On the other hand, severe ancestral stress has been shown to associate with the offspring’s blood DNA methylation pattern [25,32], suggesting that ancestral exposures could also affect the descendant through DNA methylation levels in humans. However, our study setting differs somewhat from the previous studies focusing on grandparental nutritional status, as the anthropometrics have been measured during or after puberty and not during the prepubertal slow-growth period [21–23]. Moreover, in previous studies, harvest records were utilized as a proxy for food availability, whereas the BMI of the G_A_ men born between 1904 and 1937 should not be interpreted as a straightforward indicator of food availability in the context of our study.

As regards anthropometric variables, especially adult height is understood to capture the cumulative conditions of childhood and adolescence, including food availability [33–35]. However, we identified only modest associations between the height of the G_A_ men and the epigenetic ageing of their descendants, and this association was direct, as opposed to the indirect association observed for BMI and weight. Complicating the interpretation of our results was the fact that not all of the men in our cohort had reached their adult height, as records from the period show some of them not reaching their adult height until the age of 24 years [36]. Anthropometric measures in this historical period in Finland show a complex association not only with socioeconomic status, but also with urbanization and climate [37]. In our data for G_A_, height and weight were associated with social class, with the higher classes being taller and heavier. However, in our dataset, there was no statistically significant association between social class and BMI. In a comparable cohort of Finnish men, the upper- and middle-class men had the lowest BMI, whereas farmers had the highest BMI [37]. Before the 1950s, Finland experienced recurrent food insecurity due to climate and agricultural limitations, the nutrition was monotonous and, due to a low consumption of vegetables, lacked many micronutrients. Thus, both qualitative and quantitative hunger existed in Finland during this time period [38,39], but those exposed to it cannot be easily identified.

In our internal replication, the BMI of the mother (G1) in adolescence and young adulthood was observed to associate inversely with the pace of epigenetic ageing in her children (G2) and, in a sex-stratified analysis, in her sons, but no association was observed between a father’s (G1) BMI and the epigenetic pace of ageing in his children (G2). A recent study analysed the association between parental overweight in puberty, measured using a validated figural drawing scale describing body shape, and the epigenetic ageing of offspring in young adulthood [40]. Contrary to our results, the authors showed a direct association between paternal overweight in puberty and the epigenetic ageing of offspring, with no association between maternal overweight in puberty and the epigenetic ageing of offspring. These seemingly contradicting results might reflect the complex associations between food availability and nutrition, environmental conditions in different historical contexts and differences in developmental timing, as well as sexual dimorphism in these traits. In addition, Lopez-Cervantes et al. focus on overweight as a categorical phenotype, whereas our analysis utilized BMI as a continuous range.

Other previous works studying the association between maternal weight or BMI and the epigenetic ageing of offspring have focused on young children. Only the first-generation Horvath clock has also been trained with children, and it has been shown to tick faster in childhood than in adulthood [41], which is why a comparison of these results to those obtained from adult offspring is not without issues. Fransquet et al. have reported an inverse association between maternal weight and epigenetic age deviation in the offspring as measured with both the Horvath clock and the paediatric buccal cell clock PedBE [42] when the children were approximately 1 year old. In contrast to this finding, it has also been reported that maternal BMI during pregnancy is directly associated with epigenetic age deviation in childhood, but this study also only included the Horvath clock [43].

While BMI is, to an extent, heritable, our results do not reflect the heritability of BMI and its close link to epigenetic ageing. An individual’s own increased BMI has a direct association with epigenetic ageing, with a higher BMI being associated with an accelerated DunedinPACE [44,45] and increased age deviation in other epigenetic clocks [45–48], whereas our results show an inverse association between an individual’s BMI in young adulthood and the offspring’s DunedinPACE. In addition, BMI has been shown to be more inherited in the maternal line, possibly through intrauterine conditions [49] or social heritability of eating habits [50], although the idea that parental dietary intake behaviour shapes a child’s dietary intake has also been challenged [51].

While we can show an association between grandparental anthropometric measures and descendants’ phenotypes, our data do not provide the means to establish whether we are observing transgenerational inheritance, repeated intergenerational inheritance, or a non-biological inheritance of phenotypes. By definition, grandpaternal environmental exposures leading to effects in grandchildren would be transgenerational inheritance, as the germ cells giving rise to the offspring have not been directly exposed [20]. However, a parent having a higher BMI could cause the epigenetic programming of the gametes, leading to both a higher BMI and a slower epigenetic pace of ageing in the child. This would again lead to similar reprogramming of the grandchild, resulting in repeated intergenerational inheritance. This scenario is not, however, in line with the literature which shows that an individual’s higher BMI is generally associated with a faster pace of epigenetic ageing.

Our results could also be indicative of non-biological inheritance. Epigenetic clocks can be interpreted to measure general well-being (or ill-being), as they are good predictors of morbidity and mortality [29,30]. These results could thus be describing the inheritance of socioeconomic status from the G_A_ men to their offspring through the maternal line, as a higher BMI at drafting was associated with a slower pace of epigenetic ageing in the descendants. Socioeconomic status has also been shown to be strongly inherited in Finland [52], but in general, it has been demonstrated to be more inherited by sons from fathers, with daughters presenting more social mobility [53]. Similarly, a father’s educational level has a stronger association with the educational attainment of their sons than that of their daughters [54,55]. Our analyses have also been adjusted for the socioeconomic status of the G_A_ men and the years of schooling completed by the descendants.

One final putative explanation between the grandparental or parental BMI and the pace of epigenetic ageing in the descendants could be the association of growth and development with the ageing process, as proposed by the developmental theory of ageing [56,57]. If the BMI in adolescence or young adulthood also reflects the intrinsic pace of development in addition to the environmental and lifestyle effects, the inheritance of the intrinsic pace of ageing could also be reflected in the pace of epigenetic ageing in the offspring. However, as our results show an inverse association between parental BMI and the epigenetic ageing of offspring, while Lopez-Cervantes et al. report accelerated epigenetic ageing with parental obesity, this interpretation of our results remains highly speculative.

We saw no association between the G_A_ men’s anthropometric measurements and the genome-wide methylation patterns in the blood of their descendants, with the exception of one site, where the grandpaternal weight associated with the DNA methylation levels at cg10866062 in the transcription start site of *BTBD17* in the maternal granddaughters. Similarly, in the sperm of sons and grandsons, one statistically significant differentially methylated locus that associates with the G_A_ men’s early adulthood BMI was identified for each of the two generations, both located in uncharacterized lncRNA candidate genes. These very limited findings do not support the role of DNA methylation as a molecular mediator in a transgenerational or intergenerational inheritance of paternal nutrition status. Even though our sample size is larger than in previous studies with similar settings [25,26], it is still limited for a genome-wide association analysis, potentially leading to a lack of power to identify milder effects, but we would expect that if the effects were systematic or substantial, we could see them in our multigenerational setting.

The relatively low number of individuals per generation is a clear limitation of our study, as is the lack of insight into the determinants of BMI in the G_A_ men. In addition, we were only able to connect the G_A_ men’s data to their descendants, if the men were alive in 1964, leading to potential selection bias. However, we utilized an extremely unique data series with individual measurements from the G_A_ men and more than 100 years of follow-up from their descendants, with epigenetic profiling available from three generations. In addition to utilizing DNA methylation data from blood, we also utilized a smaller set of sperm DNA methylation data, which is more relevant in conveying epigenetic information to the offspring.

In conclusion, we show that a higher BMI in the G_A_ men is associated with a slower pace of epigenetic ageing in their daughters and their daughters’ daughters. We are thus able to show the inheritance of health-related traits and their association to parental phenotypes in multiple generations. However, we were not able to find evidence for the potential mechanisms of this inheritance, as there were no associations to methylation profiles at large in either blood or sperm. Even though we measured health via epigenetic clocks and, therefore, DNA methylation, our study indicates an association of acquired traits in future generations, not epigenetic inheritance through DNA methylation patterns.

## Methods

### Cohort description

The Young Finns Study (YFS) is a longitudinal population cohort started in 1980 with participants aged between 3 and 18 years and regular data collection every 3–9 years [27,58]. The most recent follow-up in 2018–2020 also included the original participants’ (G1) parents (G0) and children (G2) [27]. In the present study, we added data on one additional preceding generation, the fathers of the G0 generation, referred to as G_A_, to the analysis. Their identity information was obtained from the Digital and Population Data Services Agency (DVV). Data were only available for men (G_A_) who were alive in 1964, when national population register keeping was initiated. The military service records of these men were obtained from the National Archives of Finland. These records contain data on anthropometrics, health, social background and war experience [59]. We identified a total of 1,429 men (G_A_), 994–1,013 of whom were under the age of 25 and had anthropometric data – i.e. height, weight, BMI and chest circumference – available. Genome-wide blood and sperm DNA methylation data were available for subsets of three generations of their descendants.

### Variable description

A three-level variable for the social class of the G_A_ men was created based on a slight modification of the HISCLASS12 classification [60]. The lower-class category includes lower-skilled workers, lower-skilled farm workers, unskilled workers, unskilled farm workers and general workers; the middle-class category includes foremen, medium-skilled workers, farmers and fishermen; while the upper-class category includes higher managers, higher professionals, lower managers, lower professionals, clerical and sales personnel, lower clerical and sales personnel, as well as students.

As there are known genetic and epigenetic differences between Eastern and Western Finns [61,62], the municipality of birth of the G_A_ men was used to classify them as Eastern (provinces of Viipuri, Mikkeli, Kuopio, Oulu, Lappi) or Western Finns (provinces of Uusimaa, Turku and Pori, Häme, Vaasa). This variable was used to adjust our models for geographic origins.

The height and weight of G0, G1 and G2 YFS participants were measured at the clinic visit, and the body mass index (kg/m^2^) was calculated. In the present study, the BMIs of G0 and G2 in the 2018 follow-up are used to adjust regression models. For G1, the measurements from 2011 and 2018 are used for the blood and sperm analyses, respectively, as these correspond to the time of methylation measurement. Additionally, BMI estimates of G1 individuals from the ages of 3 to 34 were calculated from long-term trends of BMI [31]. In brief, we utilized measurements from 1980 to 2011 to calculate the area under the curve (AUC) for BMI. First, participant-specific curves for the measurements were estimated by mixed model regression splines [63].Then, similarly to the approach of Lai et al. [64], the AUCs were evaluated for each measurement. Individual AUCs were then utilized to create estimate values for the measurements for each year.

Data on smoking habits and the highest completed education of the G0, G1 and G2 YFS participants were self-reported in a questionnaire. A two-group variable for smoking was used with the categories of ‘daily smoker’ and ‘smokes less than daily’ (also including non-smokers). The data on education were converted into a continuous variable representing years of schooling. When used for model adjustments, the answers collected in 2018 were used for G0 and G2, while those obtained in 2011 were used for G1, corresponding to the date of methylation measurement. The only exception was G2 education, where parental education level was used as a proxy for socio-economic status.

### YFS sample collection and processing for DNA methylation analysis

Venous blood samples were taken after a 4-hour fast and stored at −70°C until analysis. Blood samples for genome-wide DNA methylation measurement in YFS were collected in 2011 and 2018–2020 for G1, and the latter follow-up also included data from G0 and G2.

A detailed description of the semen collection method can be found in Pahkala et al., 2026 [27]. In brief, human semen from males aged 18 and older was collected into an 8 ml tube containing a mixture of penicillin/streptomycin and gentamicin sulphate. The samples were stored at −20°C until purification. A sperm count was performed, and the sample was transferred to 2 ml of 50% Puresperm (Nidacon) solution and centrifuged at 400 x g for 15 minutes. The sperm pellet was washed by adding an equal volume of 2X Somatic Cell Lysis Buffer, centrifuged at 3,000 x g for 15 minutes and further washed with 3 ml of phosphate-buffered saline (PBS), followed by a final centrifugation at 2,500 x g for 15 minutes. The sperm pellet was resuspended in 1 ml of PBS, and another sperm count was performed. Sample purity was assessed by staining with SPERM STAIN (Microptic), and the percentage of somatic cell contamination was calculated. The remaining sample was pelleted at 12,000 x g for 5 minutes, the supernatant was discarded, and the pellet was snap-frozen in liquid nitrogen and stored at −80°C.

### DNA isolation and methylation measurement

DNA for methylation analysis was isolated from EDTA peripheral blood samples using a Wizard^®^ Genomic DNA Purification Kit in the 2011 follow-up [65] and a Perkin Elmer Chemagic (CMG-1074) in the 2018–2020 follow-up [27]. Following bisulphite conversion, genome-wide DNA methylation levels were measured using the Illumina Infinium HumanMethylation450 BeadChip or the Illumina Infinium Methylation EPIC BeadChip version 1 for the 2011 follow-up [19] at Helmholtz Zentrum, Munich, Germany. In the 2018–2020 follow-up, DNA methylation was measured in two batches; the first was done with Illumina Infinium Methylation EPIC BeadChip version 1, whereas the second batch included both versions 1 and 2 [27].

DNA isolation on the sperm pellet was performed with a modified protocol using a Revvity Chemagic 360 robot with a DNA-tissue 96 kit (art.no: Chemagic kit CMG-723). In brief, the cells were incubated for one hour at room temperature in lysis buffer S with 50 mM of β-mercaptoethanol and 20 mg/ml of proteinase K, after which 120 µl of lysis buffer B was added. The mixture was incubated for 1 hour, and isolation was performed with a Chemagic 360 robot without adding the proteinase K and lysis buffer again. Libraries for Reduced Representation Bisulfite Sequencing (RRBS) were prepared using Nugen Ovation RRBS Methyl-Seq with TrueMethyl BS (Tecan), and the libraries were sequenced by the NovaSeq 6000 system (Illumina).

### Methylation data preprocessing

Pre-processing of DNA methylation data was performed using functions from the *minfi* [66], *SeSAMe* [67] and *meffil* [68] R packages. Data from the 2018–2020 follow-up were processed separately for each generation, batch and methylation array version. In brief, all pipelines included background correction using the noob method [69], dye-bias correction, normalization and exclusion of unreliable probes, as described previously [70]. Beta methylation levels were calculated and retained for downstream analyses. DNA methylation-based estimates of cell proportions were calculated using the Houseman method [71] through functions in *minfi,* for G1, and *meffil,* for G0 and G2.

Quality control of RRBS data was performed using FASTQC (version 0.12.0) [72], and reads were trimmed with TrimGalore (version 0.6.4) [73]. NuGen diversity adaptors were removed with a Python script supplied by NuGen “trimRRBSdiversityAdaptCustomers.py” (version 1.11). Reads were then aligned to the GRCh38 genome using Bismark (version 0.22.3) [74] with Hisat2 (version 2.2.1) [75]. The aligned files were then sorted and indexed using samtools (version 1.21) [76]. The quality control summary was concatenated using MultiQC (version 1.24) [77]. Downstream analysis was done using methylkit (version 3.19) [78]. Filtering was performed with a minimum phred quality score of 20, and every region where there was at least 10% overlap among all samples was called.

### Epigenetic age estimation

Principal-component versions [79] of PhenoAge [28] and GrimAge [29], designed to increase reliability, were computed with the methylCIPHER R package [80], while DunedinPACE [30] was calculated using the *DunedinPACE* R package. Participants with missing data in more than 20% of the required CpGs were excluded. The epigenetic age deviation (AgeDev) value was calculated for the principal-component derivations of the second-generation clocks (PhenoAge and GrimAge) by taking the residual obtained after regressing the epigenetic age on chronological age [81].

### Epigenetic age association

Multiple regression models were performed in R to determine the association between each anthropometric trait of the G_A_ men and the biological age of their descendants. In the regression models, the G_A_ men’s early-adulthood anthropometric measures were used as independent variables, while the descendants’ epigenetic age measures were used as dependent variables. Three models were applied for each setting. The basic model was adjusted for the descendant’s age, sex, BMI, smoking status, cell proportions, DNA methylation array and, for data collected in 2018, the batch. In the adjusted model, the G_A_ men’s social class, geographic origin, year of birth and lineage, when applicable, were further accounted for. The fully adjusted model was additionally adjusted for the years of schooling for children (G0) and grandchildren (G1), or the parent’s years of schooling for great-grandchildren (G2). Participant selection to maintain variable independence was performed as described for the EWAS analysis. The analyses were repeated with two binary variables comparing participants in either the upper or lower quartile of BMI to the rest.

Internal replication was performed using the G1’s estimated BMI between the ages of 3 and 34 and G2’s epigenetic age. Multiple regression models were adjusted for G2 age, smoking status and BMI; G1 age, geographic origin and years of schooling; as well as DNA methylation batch, array version and estimated cell proportions. Analyses were performed separately for mothers and fathers with all children and then stratified by offspring sex. When data from more than one G2 sibling was available, one individual was selected from each family cluster at random to maintain variable independence.

### EWAS in blood

Epigenome-wide association studies (EWASs) were performed to determine whether G_A_ anthropometrics associate with the descendants’ DNA methylation at each of the loci measured by the arrays. Analyses of the children (G0) were performed using methylation data collected in 2018 with the Illumina epic v2 array including 894,342 loci. Children were analysed all together (n = 250) and separately for sons (n = 133) and daughters (n = 117). Analyses of the grandchildren (G1) were performed using data from the 2011 collection due to a larger sample size, with the Illumina epic v1 array including 769,683 probes. Grandchildren were analysed all together (n = 392), as well as separately for paternal grandchildren (n = 242), maternal grandchildren (n = 253), paternal grandsons (n = 100) and maternal granddaughters (n = 121). Analyses of the great-grandchildren (G2) were performed using data collected in 2018 with the Illumina epic v2 array including 894,877 probes. Great-grandchildren were analysed all together (n = 143), as stratified analyses would not have had sufficient statistical power. In cases where data from more than one grandfather or great-grandfather were available, they were selected at random to avoid participant duplication. All analyses were performed using multiple regression models adjusted for the G_A_ men’s social class, geographic origin and year of birth, the descendant’s age, BMI and smoking status, as well as cell proportions and the first 10 principal components of the methylation control probes. Where appropriate, models were also adjusted for lineage and the descendant’s sex. To account for multiple testing, the Benjamini Hochberg false discovery rate was calculated, and the threshold for statistical significance was FDR < 0.05.

### EWAS in sperm

The association between the G_A_ men’s BMI and the RRBS DNA methylation data obtained from the sperm of their sons (G0, n = 51) and grandsons (G1, n = 98) was analysed using the *DSS* R package [82]. The analysis for great-grandsons (G2) could not be performed, as data were only available for 24 individuals. The analysis was performed by first fitting a beta-binomial generalized linear model at each locus and then performing a Wald test for each of them to determine significantly differential methylation. Benjamini-Hochberg correction for multiple testing was applied. Analyses were performed separately for sons (G0) and grandsons (G1) of the G_A_ men. The model coefficients included variables from the G_A_ men – age at BMI measurement, social class, geographic origin – and from the descendant – age at methylation measurement, BMI, smoking status and lineage (the latter only for grandsons).

## Declarations

### Ethics approval and consent to participate

The YFS has been approved by the 1st ethical committee of the Hospital District of Southwest Finland and by local ethical committees (1st Ethical Committee of the Hospital District of Southwest Finland, Regional Ethics Committee of the Expert Responsibility area of Tampere University Hospital, Helsinki University Hospital Ethical Committee of Medicine, Research Ethics Committee of the Northern Savo Hospital District, and the Ethics Committee of the Northern Ostrobothnia Hospital District). All participants of these studies have given informed consent, and the studies have been conducted according to the principles of the Declaration of Helsinki.

## Consent for publication

Not applicable

## Availability of data and materials

The datasets generated and/or analysed during the current study are not publicly available due to restrictions imposed by Finnish legislation but are available from the corresponding author/data sharing committees upon a reasonable request.

## Authors’ contribution

SM, IT, VL, VK and ER planned the study. IT, VL and VK acquired new data. JC, SM, IT, VL, VK and ER designed the data analysis. JC performed the analyses, while SM, NKa, PPM, DK, LPL and ER instructed in the data analysis. NKo, DK, PPM, LPL, SR, JM, KP, MK, OR, TL and ER provided data. OR, TL, VK and ER acquired funding and provided general leadership. JC and ER wrote the first draft of the manuscript and edited it in response to the comments of the co-authors. JC, SM, IT, VL, VK and ER participated in the writing of the manuscript, and all co-authors read and revised the final manuscript.

## References

1. Blake GE, Watson ED. Unravelling the complex mechanisms of transgenerational epigenetic inheritance. Current Opinion in Chemical Biology. 2016;33:101–7. 10.1016/j.cbpa.2016.06.008

2. Anway MD, Leathers C, Skinner MK. Endocrine disruptor vinclozolin induced epigenetic transgenerational adult-onset disease. Endocrinology. 2006;147. 10.1210/en.2006-0640

3. Van Steenwyk G, Roszkowski M, Manuella F, Franklin TB, Mansuy IM. Transgenerational inheritance of behavioral and metabolic effects of paternal exposure to traumatic stress in early postnatal life: Evidence in the 4th generation. Environmental Epigenetics. 2018;4. 10.1093/eep/dvy023

4. Gapp K, Jawaid A, Sarkies P, Bohacek J, Pelczar P, Prados J, et al. Implication of sperm RNAs in transgenerational inheritance of the effects of early trauma in mice. Nat Neurosci. 2014;17:667–9. 10.1038/nn.3695

5. Chen Q, Yan M, Cao Z, Li X, Zhang Y, Shi J, et al. Sperm tsRNAs contribute to intergenerational inheritance of an acquired metabolic disorder. Science. 2016;351:397–400. 10.1126/science.aad7977

6. Sharma U, Conine CC, Shea JM, Boskovic A, Derr AG, Bing XY, et al. Biogenesis and function of tRNA fragments during sperm maturation and fertilization in mammals. Science. 2016;351:391–6. 10.1126/science.aad6780

7. Grandjean V, Fourré S, De Abreu DAF, Derieppe M-A, Remy J-J, Rassoulzadegan M. RNA-mediated paternal heredity of diet-induced obesity and metabolic disorders. Sci Rep. 2015;5:18193. 10.1038/srep18193

8. Singh S, Shyam S, Sah S, Singh MK, Palta P. Treatment of buffalo (Bubalus bubalis) somatic cell nuclear transfer embryos with MicroRNA-29b mimic improves their quality, reduces DNA methylation, and changes gene expression without affecting their developmental competence. Cellular Reprogramming. 2019; 10.1089/cell.2019.0007

9. Rakyan VK, Chong S, Champ ME, Cuthbert PC, Morgan HD, Luu KVK, et al. Transgenerational inheritance of epigenetic states at the murine *Axin ^Fu^* allele occurs after maternal and paternal transmission. Proc Natl Acad Sci USA. 2003;100:2538–43. 10.1073/pnas.0436776100

10. Takahashi Y, Morales Valencia M, Yu Y, Ouchi Y, Takahashi K, Shokhirev MN, et al. Transgenerational inheritance of acquired epigenetic signatures at CpG islands in mice. Cell. 2023;186:715–731.e19. 10.1016/j.cell.2022.12.047

11. Zeng Y, Chen T. DNA methylation reprogramming during mammalian development. Genes. 2019. 10.3390/genes10040257

12. Heard E, Martienssen RA. Transgenerational epigenetic inheritance: Myths and mechanisms. Cell. 2014. 10.1016/j.cell.2014.02.045

13. Barlow DP, Bartolomei MS. Genomic imprinting in mammals. Cold Spring Harbor Perspectives in Biology. 2014;6. 10.1101/cshperspect.a018382

14. Fitz-James MH, Cavalli G. Molecular mechanisms of transgenerational epigenetic inheritance. Nat Rev Genet. Nature Publishing Group; 2022;23:325–41. 10.1038/s41576-021-00438-5

15. Jawaid A, Jehle KL, Mansuy IM. Impact of Parental Exposure on Offspring Health in Humans. Trends in Genetics. 2021. 10.1016/j.tig.2020.10.006

16. Sandin S, Hultman CM, Kolevzon A, Gross R, MacCabe JH, Reichenberg A. Advancing maternal age is associated with increasing risk for autism: A review and meta-analysis. Journal of the American Academy of Child and Adolescent Psychiatry. 2012;51. 10.1016/j.jaac.2012.02.018

17. Hultman CM, Sandin S, Levine SZ, Lichtenstein P, Reichenberg A. Advancing paternal age and risk of autism: New evidence from a population-based study and a meta-analysis of epidemiological studies. Molecular Psychiatry. 2011;16. 10.1038/mp.2010.121

18. Carpenter BL, Zhou W, Madaj Z, DeWitt AK, Ross JP, Grønbæk K, et al. Mother–child transmission of epigenetic information by tunable polymorphic imprinting. Proceedings of the National Academy of Sciences of the United States of America. 2018; 10.1073/pnas.1815005115

19. Marttila S, Viiri LE, Mishra PP, Kühnel B, Matias-Garcia PR, Lyytikäinen L-P, et al. Methylation status of nc886 epiallele reflects periconceptional conditions and is associated with glucose metabolism through nc886 RNAs. Clin Epigenet. 2021;13:143. 10.1186/s13148-021-01132-3

20. Skinner MK, Nilsson EE. Epigenetic Transgenerational Inheritance. Encyclopedia of Reproduction [Internet]. Elsevier; 2018 [cited 2026 Oct 14]. p. 436–8. 10.1016/B978-0-12-801238-3.64510-5

21. Pembrey M, Saffery R, Bygren LO lov. Human transgenerational responses to early-life experience: potential impact on development, health and biomedical research. Journal of medical genetics. 2014. 10.1136/jmedgenet-2014-102577

22. Vågerö D, Pinger PR, Aronsson V, van den Berg GJ. Paternal grandfather’s access to food predicts all-cause and cancer mortality in grandsons. Nature Communications. 2018;9. 10.1038/s41467-018-07617-9

23. van den Berg GJ, Pinger PR. Transgenerational effects of childhood conditions on third generation health and education outcomes. Economics and Human Biology. 2016;23. 10.1016/j.ehb.2016.07.001

24. Sari E, Moilanen M, Sommerseth HL. Transgenerational health effects of in utero exposure to economic hardship: Evidence from preindustrial Southern Norway. Economics & Human Biology. 2021;43:101060. 10.1016/j.ehb.2021.101060

25. Mulligan CJ, Quinn EB, Hamadmad D, Dutton CL, Nevell L, Binder AM, et al. Epigenetic signatures of intergenerational exposure to violence in three generations of Syrian refugees. Sci Rep. Nature Publishing Group; 2025;15:5945. 10.1038/s41598-025-89818-z

26. Oren G, Shoshani A, Samra NN, Verbeke WJ, Vrticka P, Aisnberg-Shafran D, et al. From trauma to resilience: psychological and epigenetic adaptations in the third generation of holocaust survivors. Sci Rep. Nature Publishing Group; 2025;15:26193. 10.1038/s41598-025-12085-5

27. Pahkala K, Rovio S, Kartiosuo N, Auranen K, Bourgery M, Elovainio M, et al. Cohort Profile Update: Expanding the Cardiovascular Risk in Young Finns Study into a multigenerational cohort. International Journal of Epidemiology. 2026;55:dyaf206. 10.1093/ije/dyaf206

28. Levine ME, Lu AT, Quach A, Chen BH, Assimes TL, Bandinelli S, et al. An epigenetic biomarker of aging for lifespan and healthspan. Aging. 2018;10:573–91. 10.18632/aging.101414

29. Lu AT, Quach A, Wilson JG, Reiner AP, Aviv A, Raj K, et al. DNA methylation GrimAge strongly predicts lifespan and healthspan. Aging. 2019;11:303–27. 10.18632/aging.101684

30. Belsky DW, Caspi A, Corcoran DL, Sugden K, Poulton R, Arseneault L, et al. DunedinPACE, a DNA methylation biomarker of the pace of aging. eLife. 2022;11:e73420. 10.7554/eLife.73420

31. Koskinen JS, Kytö V, Juonala M, Viikari JSA, Nevalainen J, Kähönen M, et al. Childhood risk factors and carotid atherosclerotic plaque in adulthood: The Cardiovascular Risk in Young Finns Study. Atherosclerosis. 2020; 10.1016/j.atherosclerosis.2019.11.029

32. Yehuda R, Daskalakis NP, Bierer LM, Bader HN, Klengel T, Holsboer F, et al. Holocaust Exposure Induced Intergenerational Effects on FKBP5 Methylation. Biol Psychiatry. 2016;80:372–80. 10.1016/j.biopsych.2015.08.005

33. Floud R, National Bureau of Economic Research, editors. The changing body: health, nutrition, and human development in the western world since 1700. Cambridge; New York: Cambridge University Press; 2011.

34. Harris B. Anthropometric history and the measurement of wellbeing. Populationyearbook. 2021;19:91. 10.1553/populationyearbook2021.rev02

35. Steckel RH. Heights and human welfare: Recent developments and new directions. Explorations in Economic History. 2009;46:1–23. 10.1016/j.eeh.2008.12.001

36. Lindén VF. Asevelvollistemme kuntoisuus ja kansanravitsemus. Helsinki;

37. Quanjer B, Kruit JF, Taskinen I. Beyond cold and class: drivers of anthropometric differences in modernising Finland. Scandinavian Economic History Review. 2026;1–34. 10.1080/03585522.2026.2721255

38. Kovalainen N, Niemi J, Huan-Niemi E, 4100310210, 4100310210, Luonnonvarakeskus. Review of food security in Finland from the 17th to 21st century [Internet]. Luonnonvarakeskus; 2024 [cited 2026 May 21]. https://jukuri.luke.fi/handle/11111/9625. Accessed 21 May 2026

39. Hjelt V. Tutkimus ammattityöläisten toimeentuloehdoista Suomessa 1908-1909. Helsinki: Työtilasto XIII; 1912.

40. López-Cervantes JP, Mørkve Østergaard T, Kitaba NT, Lønnebotn M, Bertelsen RJ, Accordini S, et al. Epigenetic age acceleration in offspring linked to paternal smoking initiation and overweight in puberty: Evidence from a two-generation study. Environmental Advances. 2026;25:100739. 10.1016/j.envadv.2026.100739

41. Horvath S. Erratum to: DNA methylation age of human tissues and cell types. Genome Biol. 2015;16:96. 10.1186/s13059-015-0649-6

42. Fransquet PD, Macdonald JA, Ryan J, Greenwood CJ, Olsson CA. Exploring Perinatal Biopsychosocial Factors and Epigenetic Age in 1-Year-Old Offspring. Epigenomics. Taylor & Francis; 2023;15:927–39. 10.2217/epi-2023-0284

43. Simpkin AJ, Hemani G, Suderman M, Gaunt TR, Lyttleton O, Mcardle WL, et al. Prenatal and early life influences on epigenetic age in children: a study of mother-offspring pairs from two cohort studies. Hum Mol Genet. 2016;25:191–201. 10.1093/hmg/ddv456

44. Drouard G, Argentieri MA, Heikkinen A, Ollikainen M, Kaprio J. Associations Between 40-Year Trajectories of BMI and Proteomic and Epigenetic Aging Clocks: Deciphering Nonlinearity and Interactions. Aging Cell. 2026;25:e70397. 10.1111/acel.70397

45. Li DL, Hodge AM, Cribb L, Southey MC, Giles GG, Milne RL, et al. Body Size, Diet Quality, and Epigenetic Aging: Cross-Sectional and Longitudinal Analyses. J Gerontol A Biol Sci Med Sci. 2024;79:glae026. 10.1093/gerona/glae026

46. Lundgren S, Kuitunen S, Pietiläinen KH, Hurme M, Kähönen M, Männistö S, et al. BMI is positively associated with accelerated epigenetic aging in twin pairs discordant for body mass index. Journal of Internal Medicine. 2022;292:627–40. 10.1111/joim.13528

47. Cao X, Yang G, Li X, Fu J, Mohedaner M, Danzengzhuoga, et al. Weight change across adulthood and accelerated biological aging in middle-aged and older adults. The American Journal of Clinical Nutrition. 2023;117:1–11. 10.1016/j.ajcnut.2022.10.020

48. Ho KM. Associations between body mass index, biological age and frailty in the critically ill. Obesity Research & Clinical Practice. 2024;18:189–94. 10.1016/j.orcp.2024.05.004

49. Kelly GE, Murrin C, Viljoen K, O’Brien J, Kelleher C. Body mass index is associated with the maternal lines but height is heritable across family lines in the Lifeways Cross-Generation Cohort Study. BMJ Open. 2014;4:e005732. 10.1136/bmjopen-2014-005732

50. Akbarzadeh M, Teymoori F, Riahi P, Farhadnejad H, Ahmadirad H, Sadat Zahedi A, et al. Evidence of familial resemblance and family-based heritability of food intakes derived from a longitudinal cohort study. Sci Rep. Nature Publishing Group; 2023;13:11934. 10.1038/s41598-023-38326-z

51. Pervin S, Emmett P, Townsend N, Biswas T, Huda MM, Northstone K, et al. The myth and reality of familial resemblance in dietary intake: a systematic review and meta-analysis on the resemblance of dietary intake among parent and offspring. eClinicalMedicine. 2023;60:102024. 10.1016/j.eclinm.2023.102024

52. Lucas REB, Kerr SP. Intergenerational income immobility in Finland: contrasting roles for parental earnings and family income. Journal of Population Economics. Springer; 2013;26:1057–94.

53. Hirvonen LH. Intergenerational Earnings Mobility Among Daughters and Sons: Evidence from Sweden and a Comparison with the United States. The American Journal of Economics and Sociology. 2008;67:777–826. 10.1111/j.1536-7150.2008.00598.x

54. Hautala H, Kallio JM. Gender matters: family background and upper secondary education in Finland. Journal of Poverty and Social Justice. Policy Press; 2021;29:47–65. 10.1332/175982720X16040856541379

55. Schneebaum A, Rumplmaier B, Altzinger W. Intergenerational Educational Persistence in Europe [Internet]. Vienna University of Economics and Business; 2014 [cited 2026 Sept 30]. 10.57938/17137FF0-E4A9-40BB-9726-E671E4AF1BD5

56. De Magalhães JP. Programmatic features of aging originating in development: aging mechanisms beyond molecular damage? The FASEB Journal. 2012;26:4821–6. 10.1096/fj.12-210872

57. Gems D. The hyperfunction theory: An emerging paradigm for the biology of aging. Ageing Research Reviews. 2022;74:101557. 10.1016/j.arr.2021.101557

58. Raitakari OT, Juonala M, Ronnemaa T, Keltikangas-Jarvinen L, Rasanen L, Pietikainen M, et al. Cohort Profile: The Cardiovascular Risk in Young Finns Study. International Journal of Epidemiology. 2008;37:1220–6. 10.1093/ije/dym225

59. Taskinen I. Construction of the Finnish Army in World War II Database. Historical Life Course Studies. 2023;13:44–60. 10.51964/hlcs13565

60. Tammes P. HISCLASS: A Historical International Social Class Scheme. By Marco H. D. Van Leeuwen and Ineke Maas (Leuven, Leuven University Press, 2011)184 pp. $49.00. The Journal of Interdisciplinary History. 2012;43:467–8. 10.1162/JINH_r_00430

61. Salmela E, Lappalainen T, Fransson I, Andersen PM, Dahlman-Wright K, Fiebig A, et al. Genome-Wide Analysis of Single Nucleotide Polymorphisms Uncovers Population Structure in Northern Europe. Gemmell NJ, editor. PLoS ONE. 2008;3:e3519. 10.1371/journal.pone.0003519

62. Ciantar J, Marttila S, Rajić S, Kostiniuk D, Mishra PP, Lyytikäinen L-P, et al. Identification and functional characterisation of DNA methylation differences between East- and West-originating Finns. Epigenetics. 2024;19:2397297. 10.1080/15592294.2024.2397297

63. Welham SJ. Smoothing spline models for longitudinal data. Longitudinal Data Analysis. 2008. 10.1201/9781420011579.ch11

64. Lai CC, Sun D, Cen R, Wang J, Li S, Fernandez-Alonso C, et al. Impact of long-term burden of excessive adiposity and elevated blood pressure from childhood on adulthood left ventricular remodeling patterns: The bogalusa heart study. Journal of the American College of Cardiology. 2014; 10.1016/j.jacc.2014.05.072

65. Mishra PP, Mishra BH, Raitoharju E, Mononen N, Viikari J, Juonala M, et al. Gene Set Based Integrated Methylome and Transcriptome Analysis Reveals Potential Molecular Mechanisms Linking Cigarette Smoking and Related Diseases. OMICS: A Journal of Integrative Biology. 2023;27:193–204. 10.1089/omi.2023.0028

66. Fortin J-P, Triche TJ, Hansen KD. Preprocessing, normalization and integration of the Illumina HumanMethylationEPIC array with minfi. Hancock J, editor. Bioinformatics. 2017;33:558–60. 10.1093/bioinformatics/btw691

67. Zhou W, Triche TJ, Laird PW, Shen H. SeSAMe: reducing artifactual detection of DNA methylation by Infinium BeadChips in genomic deletions. Nucleic Acids Research [Internet]. 2018 [cited 2026 Sept 30]; 10.1093/nar/gky691

68. Min JL, Hemani G, Davey Smith G, Relton C, Suderman M. Meffil: efficient normalization and analysis of very large DNA methylation datasets. Hancock J, editor. Bioinformatics. 2018;34:3983–9. 10.1093/bioinformatics/bty476

69. Triche TJ, Weisenberger DJ, Van Den Berg D, Laird PW, Siegmund KD. Low-level processing of Illumina Infinium DNA Methylation BeadArrays. Nucleic Acids Research. 2013;41:e90–e90. 10.1093/nar/gkt090

70. Ciantar J, Rajić S, Kostiniuk D, Raulamo E, Kartiosuo N, Lai L, et al. Genome-wide DNA methylation patterns for indicators of liver steatosis: a longitudinal multiomic study. Clin Epigenet. 2026;18:22. 10.1186/s13148-025-02037-1

71. Houseman EA, Accomando WP, Koestler DC, Christensen BC, Marsit CJ, Nelson HH, et al. DNA methylation arrays as surrogate measures of cell mixture distribution. BMC Bioinformatics. 2012;13:86. 10.1186/1471-2105-13-86

72. Andrews S. FastQC: A Quality Control Tool for High Throughput Sequence Data. http://www.bioinformatics.babraham.ac.uk/projects/fastqc/

73. Krueger F, Ewels P, He D. FelixKrueger/TrimGalore: v2.2.0 [Internet]. Zenodo; 2026 [cited 2026 June 5]. 10.5281/ZENODO.20127893

74. Krueger F, Andrews SR. Bismark: a flexible aligner and methylation caller for Bisulfite-Seq applications. Bioinformatics. 2011;27:1571–2. 10.1093/bioinformatics/btr167

75. Kim D, Paggi JM, Park C, Bennett C, Salzberg SL. Graph-based genome alignment and genotyping with HISAT2 and HISAT-genotype. Nat Biotechnol. 2019;37:907–15. 10.1038/s41587-019-0201-4

76. Li H, Handsaker B, Wysoker A, Fennell T, Ruan J, Homer N, et al. The Sequence Alignment/Map format and SAMtools. Bioinformatics. 2009;25:2078–9. 10.1093/bioinformatics/btp352

77. Ewels P, Magnusson M, Lundin S, Käller M. MultiQC: summarize analysis results for multiple tools and samples in a single report. Bioinformatics. 2016;32:3047–8. 10.1093/bioinformatics/btw354

78. Akalin A, Kormaksson M, Li S, Garrett-Bakelman FE, Figueroa ME, Melnick A, et al. methylKit: a comprehensive R package for the analysis of genome-wide DNA methylation profiles. Genome Biol. 2012;13:R87. 10.1186/gb-2012-13-10-r87

79. Higgins-Chen AT, Thrush KL, Wang Y, Minteer CJ, Kuo P-L, Wang M, et al. A computational solution for bolstering reliability of epigenetic clocks: Implications for clinical trials and longitudinal tracking. Nat Aging. 2022;2:644–61. 10.1038/s43587-022-00248-2

80. Thrush KL, Higgins-Chen AT, Liu Z, Levine ME. R methylCIPHER: A Methylation Clock Investigational Package for Hypothesis-Driven Evaluation & Research [Internet]. Bioinformatics; 2022 [cited 2026 Sept 30]. 10.1101/2022.07.13.499978

81. Chen BH, Marioni RE, Colicino E, Peters MJ, Ward-Caviness CK, Tsai P-C, et al. DNA methylation-based measures of biological age: meta-analysis predicting time to death. Aging. 2016;8:1844–65. 10.18632/aging.101020

82. Park Y, Wu H. Differential methylation analysis for BS-seq data under general experimental design. Bioinformatics. 2016;32:1446–53. 10.1093/bioinformatics/btw026

