## Supplementary Figures for "Male youth BMI is associated with epigenetic ageing of female descendants but not blood or sperm DNA methylation"

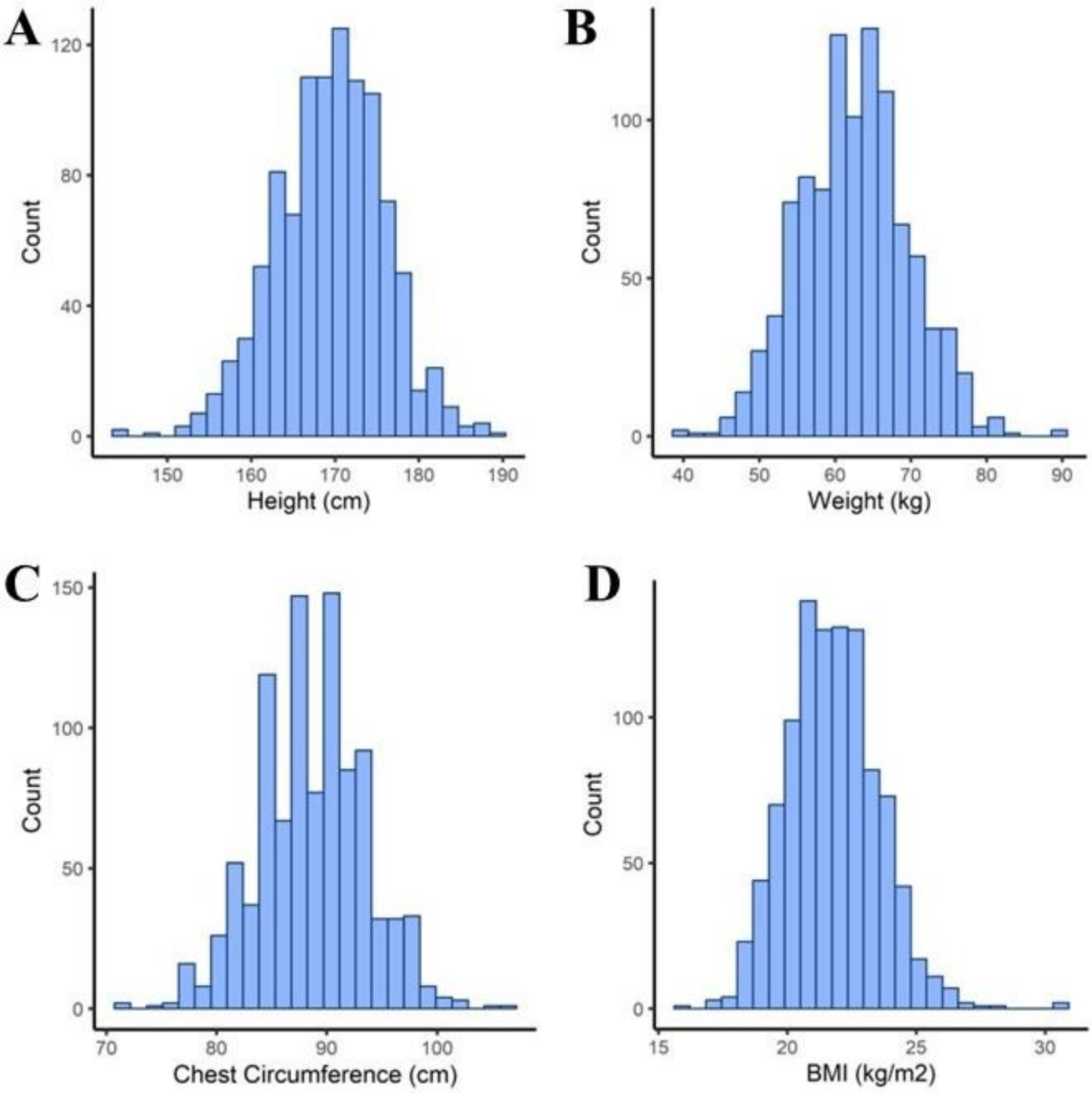


**Supplementary Figure 1**: Distribution of $G_A$ men's anthropometric measurements taken during military drafting (at ages 18-24)

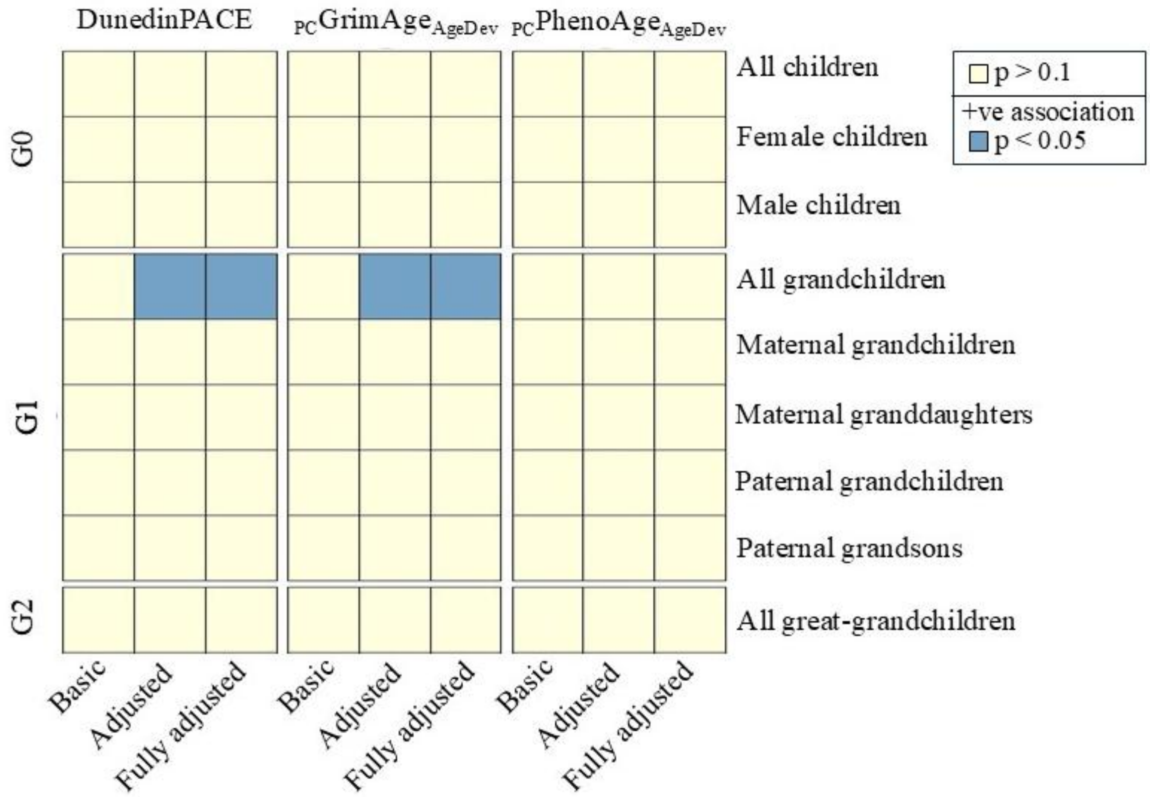


**Supplementary Figure 2**: The association between $G_A$ men's height and their descendants' epigenetic age. The blue panels represent a direct association ($p < 0.05$). The basic model was adjusted for the descendant's age, sex, BMI, smoking status, methylation-based estimates of cell proportions, as well as the DNA methylation array and batch, when applicable. The adjusted model was further adjusted for the $G_A$ men's social class, geographic origin, year of birth and lineage, when applicable. The fully adjusted model was additionally adjusted for the years of schooling that the descendant or, in the case of G2, their parent had completed.

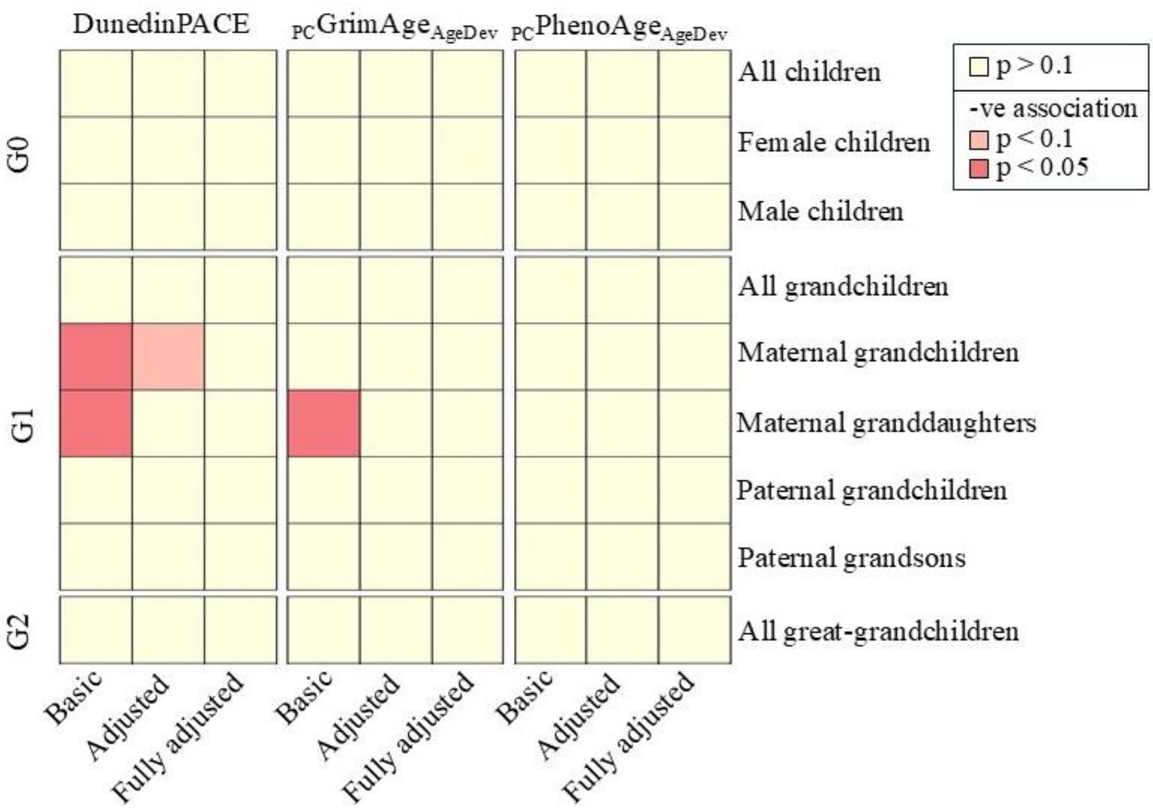


**Supplementary Figure 3**: The association between $G_A$ men's weight and their descendants' epigenetic age. The red panels represent an indirect association with the colour intensity corresponding to p value. The basic model was adjusted for the descendant's age, sex, BMI, smoking status, methylation-based estimates of cell proportions, as well as the DNA methylation array and batch, when applicable. The adjusted model was further adjusted for the $G_A$ men's social class, geographic origin, year of birth and lineage, when applicable. The fully adjusted model was additionally adjusted for the years of schooling that the descendant or, in the case of G2, their parent had completed.

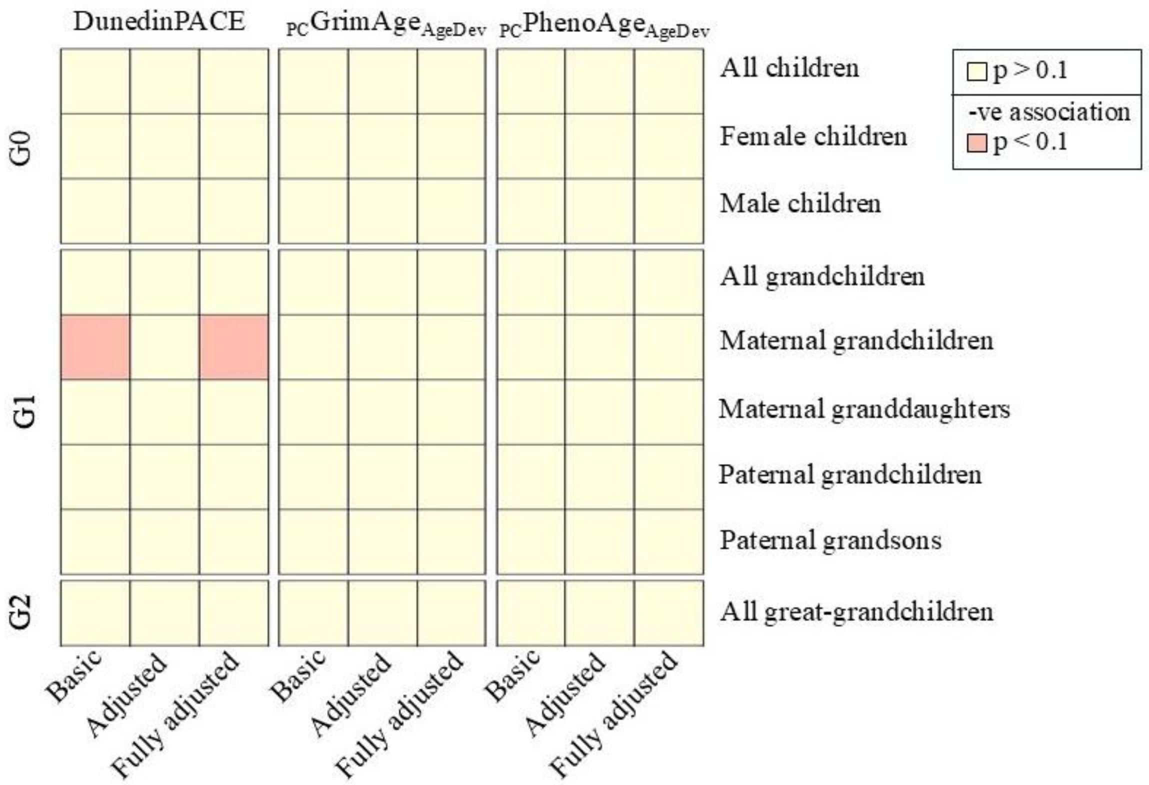


**Supplementary Figure 4**: The association between $G_A$ men's chest circumference and their descendants' epigenetic age. The light red panels represent suggest a weak indirect association ($p < 0.1$). The basic model was adjusted for the descendant's age, sex, BMI, smoking status, methylation-based estimates of cell proportions, as well as the DNA methylation array and batch, when applicable. The adjusted model was further adjusted for the $G_A$ men's social class, geographic origin, year of birth and lineage, when applicable. The fully adjusted model was additionally adjusted for the years of schooling that the descendant or, in the case of G2, their parent had completed.
